# Sucrose-mediated formation and adhesion strength of *Streptococcus mutans* biofilms on titanium

**DOI:** 10.1101/2022.09.08.507119

**Authors:** Laura J. Waldman, Tony Butera, James D. Boyd, Martha E. Grady

## Abstract

Biofilms consist of bacterial cells surrounded by a matrix of extracellular polymeric substance (EPS), which protects the colony from many countermeasures, including antibiotic treatments. Biofilm EPS composition is affected by environmental factors. In the oral cavity, the presence of sucrose affects the growth of *Streptococcus mutans* that produce acids, eroding enamel and forming dental caries. Biofilm formation on dental implants commonly leads to severe infections and failure of the implant. This work determines the effect of sucrose concentration on biofilm EPS formation and adhesion of *Streptococcus mutans*, a common oral colonizer. Bacterial biofilms are grown with varying concentrations of sucrose on titanium substrates simulating dental implant material. Strategies for measuring adhesion for films such as peel tests are inadequate for biofilms, which have low cohesive strength and will fall apart when tensile loading is applied directly. The laser spallation technique is used to apply stress wave loading to the biofilm, causing the biofilm to delaminate at a critical tensile stress threshold. Biofilm formation and EPS structures are visualized at high magnification with scanning electron microscopy (SEM). Biofilm substrate coverage and adhesion strength of biofilms initially increase with increasing sucrose concentration, but then decrease as sucrose concentration continues to increase. For biofilms grown with non-zero concentrations of sucrose, *S. mutans* adhesion to the substrate is higher than the adhesion of osteoblast-like cells to the same substrates. These results suggest sucrose-mediated adhesion and formation on titanium of *S. mutans* biofilms may outcompete osteoblasts during osseointegration, which could explain higher rates of peri-implant disease associated with high sugar diets.

## 1. INTRODUCTION

Bacterial infections are a recurring concern for permanent structural implants such as hip and dental implants, as well as temporary implants such as urinary tract devices. The high rate of infections is due in part to the way that bacterial biofilms form on implant surfaces. Bacterial cells in a biofilm are surrounded by extracellular polymeric substance (EPS) that protects the colony from treatment strategies such as antibiotics or mechanical removal [1-3]. The EPS matrix contributes to the cohesion between individual bacterial cells, as well as adhesion of the biofilm to the substrate surface [4].

Additionally, biofilm growth is affected by specific environmental factors in the growth area, including substrate material, surface roughness, hydrophobicity, and hydrodynamic pressure and temperature. Other factors known to affect biofilm formation include the concentration of nutrients such as sucrose, glucose, and other carbohydrates, as well as bacterial age and species present within the biofilm [5-10]. *Streptococcus mutans* is a common oral colonizer that causes dental caries [11] and is often found in the multispecies biofilms that grow on dental implants and can contribute to long term health issues surrounding implants [12-14]. Previous studies have demonstrated that increasing sucrose concentration increases the presence of *S. mutans* and the amount of EPS in the biofilm, resulting in thicker biofilms [11, 15]. Although these studies provide information on the constituents of the biofilm, the correlation between the sucrose concentration and the adhesion strength of the bulk biofilm to the substrate is not yet experimentally determined.

Environmental factors also impact how biofilms adhere to the substrate surface, but biofilm adhesion characterization is challenging due to the complexity of the material. Because biofilms are heterogenous and exhibit low cohesive strength, many of the methods used for thin film adhesion, such as pull tests, are incompatible with biofilms [6]. Some methods examine the interactions between a single bacterial cell and the surface using atomic force microscopy, but these techniques are not applicable to the bulk biofilm [16]. Other methods for quantifying biofilm formation examine cell presence or EPS presence within the biofilm but do not provide force-based information on the interaction between the biofilm and the substrate surface [17].

The laser spallation technique is a quantitative adhesion strength measurement method that applies stress wave loading, causing separation of a film of interest from the substrate at a critical stress value [18-20]. Because the stress load is applied indirectly to the material interface through impingement of a pulsed laser on the back side of the specimen, laser spallation is valuable for characterizing bulk adhesion properties of a wide range of materials. Material interfaces as diverse as gels [21], laminated composites [22], thin metal films [23, 24], polymer films [25], and mammalian cells on substrates [26] have been probed using the laser spallation technique. The shock loading conditions are also used to quantify dynamic behaviors of materials [27, 28] and in the development of stress-reporting materials [29]. The laser spallation technique is advantageous in cases where the cohesiveness of the film is low relative to the adhesion of the film to the substrate and traditional methods like pull tests fail to separate the interface of interest.

This work examines the effects of sucrose concentration on *S. mutans* biofilm structure and formation as well as biofilm adhesion to a titanium substrate similar to those used in dental implants. Preserved biofilm structures are imaged with electron microscopy to identify changes in biofilm morphology with increased sucrose concentration. Electron microscopy combined with an EPS preservation technique provides the high resolution needed to visualize both individual cells and EPS structures within the biofilms [17, 30]. The laser spallation technique is used to quantitatively determine the effect of sucrose concentration on the adhesion strength of the biofilm to the titanium substrate.

## 2. EXPERIMENTAL

### 2.1 Bacterial Culture & Biofilm Preparation

*Streptococcus mutans* (Wild type Xc) [31] suspended in Todd Hewitt Yeast broth (THY, VWR) with 20% glycerol is kept as frozen stock in a -80 °C freezer. Frozen bacteria stock is thawed gently using a warming bath at 37°C. A 15 mL centrifuge tube was filled with 5 mL of THY and inoculated with 1 μL of bacteria stock. The inoculated centrifuge tube is placed into a warming bath at 37 °C and cultured for 24 hr. After this period, the optical density (OD) at 600 nm is measured using a GENESYS™ 30 Visible Spectrophotometer (Thermo Scientific). THY is added to each centrifuge tube to adjust the optical density of the bacterial solution to 0.7.

Biofilms are grown on substrate assemblies for laser spallation or titanium coated slides in Petri dishes for scanning electron microscopy. Substrate assemblies or titanium coated slides are sterilized using 70% ethanol in DI water followed by UV irradiation for 30 minutes. Following sterilization, 1 mL of bacterial solution with OD_600_ equal to 0.7 and 3 mL of THY plus sucrose (VWR) at chosen concentrations is added to each dish. Concentrations of sucrose in THY used in this study are: 0 mM, 37.5 mM, 75 mM, 375 mM and 750 mM sucrose. Dishes are placed in an incubator at 37 °C with 5% CO_2_ and cultured for 24 hrs. Media is aspirated and biofilms are gently rinsed with phosphate buffered saline (PBS, VWR) to remove any detached bacteria.

### 2.2 Sample Fixation

*S. mutans* biofilms are fixed with Methacarn (Methanol – 60%, Chloroform – 30%, Acetic acid (glacial) – 10% (all VWR)) [32] for 1 hr. The samples are rinsed with PBS followed by progressive dehydration in increasing ethanol (VWR) concentrations (30% for 15 min, 50% for 15 min, 70% for 15 min, 90% for 20 min, 100% for 20 min). Samples are dried in a Leica CPD 300 critical point dryer with a total cycle time of 4 hr. Samples are sputter coated with gold/palladium to prepare for electron microscopy.

### 2.3 Scanning Electron Microscopy

Samples are imaged with a FEI Quanta 250 SEM. During top-down scanning electron microscopy, 6 biofilms prepared with each of 5 sucrose concentration are imaged. Each of the prepared 30 biofilms is imaged at 4 distinct locations with increasing magnification and an accelerating voltage of 5 keV. The 5 magnifications chosen are 250x, 2500x, 5000x, 10000x, and 15000x. A total of 120 SEM images are recorded for each sucrose concentration, for a total of 600 top-down images in this study.

In addition to top-down imaging, biofilms are also imaged in profile to obtain information about biofilm formation and film thickness. Substrates with dried and sputter coated biofilms are held sideways in metal mounting springs on specimen mounts and held in place with copper tape. The edge of the biofilm and elevated features in the biofilm are imaged with the same scanning electron microscope conditions. Biofilm images are captured at a minimum of six locations along the length of one substrate for each sucrose concentration. Sequential images are captured by first focusing on the edge of the substrate and then changing the focus to image elevated biofilm features in increasing distance away from the substrate edge.

### 2.4 Adhesion Testing

The laser spallation technique is used to determine the effect of sucrose concentration on bacterial biofilm adhesion. A schematic of the experimental setup used for biofilm-substrate adhesion measurements is shown in **Figure 1**. A 1064 nm wavelength single pulsed Nd:YAG laser (Quanta-Ray INDI, Spectra-Physics) with a pulse duration of 10 ns and adjustable energy from 0 to 300 mJ is used to initiate film spallation. A laser pulse is focused to a 2.2 mm spot size and reflects vertically to impinge upon the substrate assembly.

**Figure 1:**
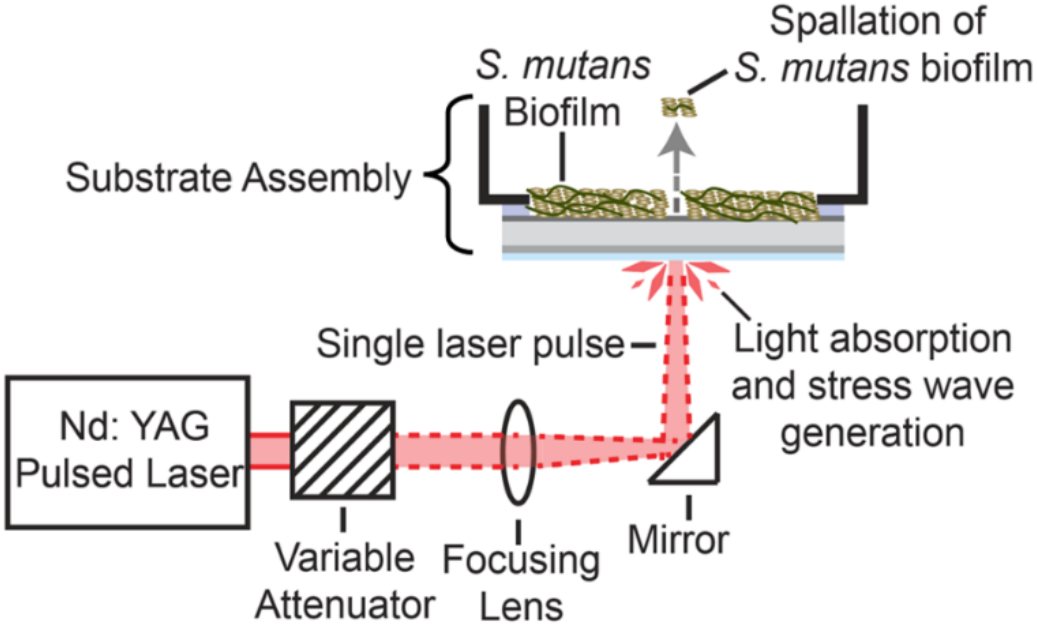
Schematic of laser spallation setup used during experimentation where a single pulse from an Nd: YAG laser impinges upon a substrate assembly resulting in ejection of *S. mutans* biofilm. Figure adapted from [19].

The substrate assembly includes a titanium-coated glass slide adhered to a 35 mm Petri dish with a 25 mm diameter hole such that a biofilm can be cultured directly on the titanium surface. On the side opposite of biofilm growth are two layers designed to generate a stress wave of sufficient amplitude to spall the biofilm. The aluminum layer (approximately 300 nm in thickness) is called the absorbing layer because it absorbs the infrared laser pulse. Because the absorbing layer is sandwiched between the substrate and a confining layer of sodium silicate (approximately 5.5 μm in thickness), the rapid gasification of the aluminum sends a large amplitude compressive wave into the substrate. The compressive stress wave reflects at the film free surface causing a tensile load on the biomaterial-titanium interface. When the amplitude of the tensile load is greater than the adhesive strength of the film, the film separates from the substrate. Each biofilm is loaded in this manner at multiple locations by adjusting translation stages. Loading regions are separated by a gap equal to the spot size to mitigate any effect from previous loads on current loading areas. The substrate assembly and the experimental method of spallation testing are discussed in greater detail in Boyd *et al*. [18, 19].

During spallation testing, *S. mutans* biofilms are loaded over a range of fluences (7.93-79.4 mJ/mm^2^), which is energy of a single pulse divided by the spot size area. Calibration experiments are performed to convert fluence values to interface stress values based on one dimensional wave propagation and transmission coefficients. Procedures for calibration experiments can be read in further detail in previous experiments [19, 20].

Spallation testing includes 12-15 loading locations per substrate assembly to determine the fluence at which failure occurs. For each sucrose concentration, 12 biofilms are tested for a total of over 100 loaded regions. Failure is recorded when visible concentric ejection of the film is observed at the loaded region. The failure rate at each fluence is used as input to a Weibull failure model to determine the adhesion strength for each biofilm.

### 2.5 Weibull Modelling of Film Failure

Due to the natural heterogeneity of biological materials, spallation occurs over a range of loading values. The failure statistics, F(σ_int,peak_), are fit to a two parameter cumulative Weibull distribution function (Equation 1) [33] to determine the adhesion strength for each sucrose concentration in this study. Weibull analysis, common in macroscopic adhesion analyses [34, 35], calculates the half-life from a Weibull distribution and is used as the adhesion strength.

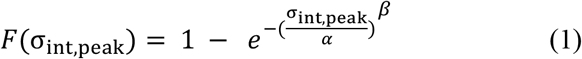

A statistical method was developed in Boyd *et al*. [19], and improved in this paper, which considers the variability in both film failure data and calibrated interface stress. Previous statistical analysis resulted in low RMS difference between the experimental film failure data and the Weibull model for *S. mutans* biofilms, resulting in asymptotic confidence intervals that were unrealistically small. Interface stress and film failure data are resampled with replacement (bootstrapped) simultaneously 1000 times to obtain simulated Weibull parameters alpha and beta. The non-linearity of the sampled beta values prevented a simple confidence interval approach. Instead, the beta values were log transformed before obtaining a 95% confidence interval, and then transformed back to the original scale. The Weibull model is interpolated for each of the 1000 simulations to obtain the interface stress that correlates to 50% failure, which represents the adhesion strength of the biofilm to the substrate. The 95% C.I. represent the range of plausible values for the median value for the adhesion strength of the biofilm to the titanium substrate.

## 3. RESULTS AND DISCUSSION

### 3.1 Optical Visualization

Stable biofilms are observed for each sucrose concentration indicated by the opaque film that forms on the titanium coated substrates as shown in **Figure 2**(b, c). Prior to bacteria deposition, the titanium surface of the substrate is clear and reflective (**Figure 2**(a)). Biofilms grown with 0 mM sucrose have incomplete coverage of the substrates, and the titanium surface can be seen through the biofilm, as shown in **Figure 2**(b). Substrates with biofilms grown with any non-zero sucrose concentration in this study, such as the 37.5 mM concentration shown in **Figure 2**(c), have more consistent coverage. For these concentrations, a cloudy and continuous film covers the substrate completely. Among biofilms grown with sucrose, visual inspection reveals very little that differentiates the samples with differing concentrations of sucrose.

**Figure 2:**
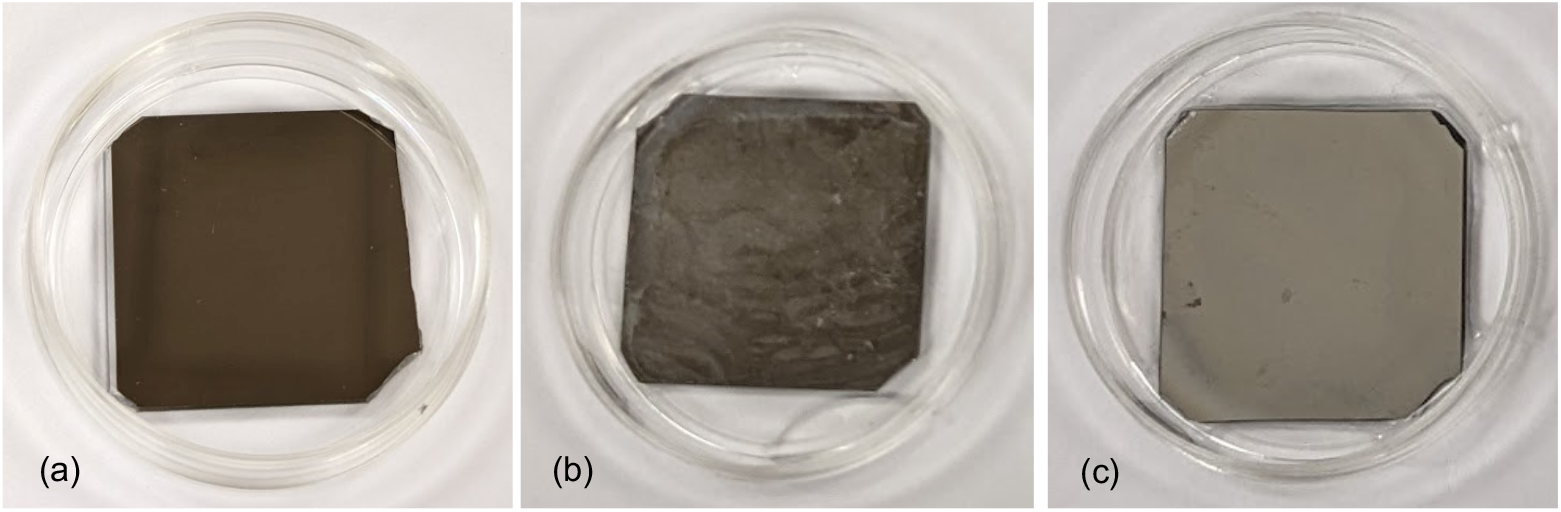
Optical images of titanium coated substrates inside 35 mm Petri dishes. (a) Bare substrate (b) Substrate with *S. mutans* biofilm grown with 0 mM sucrose (c) Substrate with *S. mutans* biofilm grown with 37.5 mM sucrose.

### 3.2 Top-Down Electron Microscopy

Biofilms are imaged at higher magnification with electron microscopy to compare biofilm structures for each sucrose concentration. At all concentrations, *S. mutans* cells form in chains and are surrounded by EPS, though the length of the chains and amount of EPS varies by sucrose concentration. Substrate coverage is not uniform; instead, the biofilms show regions with higher quantities of cells and regions that appear to be elevated. Top-down imaging reveals differences between biofilms grown with different sucrose concentrations. Biofilm samples grown in sucrose-free media form shorter chains than those with higher concentrations of sucrose and also contain very little EPS. Regions of EPS are observed in only 35% of the top-down images of biofilms, and, where present, are sparse, disconnected, and less than 5 μm in length. Though small clusters of bacteria occur on those substrates, the cells are not numerous enough to completely cover the titanium surface, which can be seen behind the cells (**Figure 3**(a)). The incomplete coverage seen in the SEM images of 0 mM sucrose biofilms agrees with the optical images of the same concentration in **Figure 2**.

**Figure 3:**
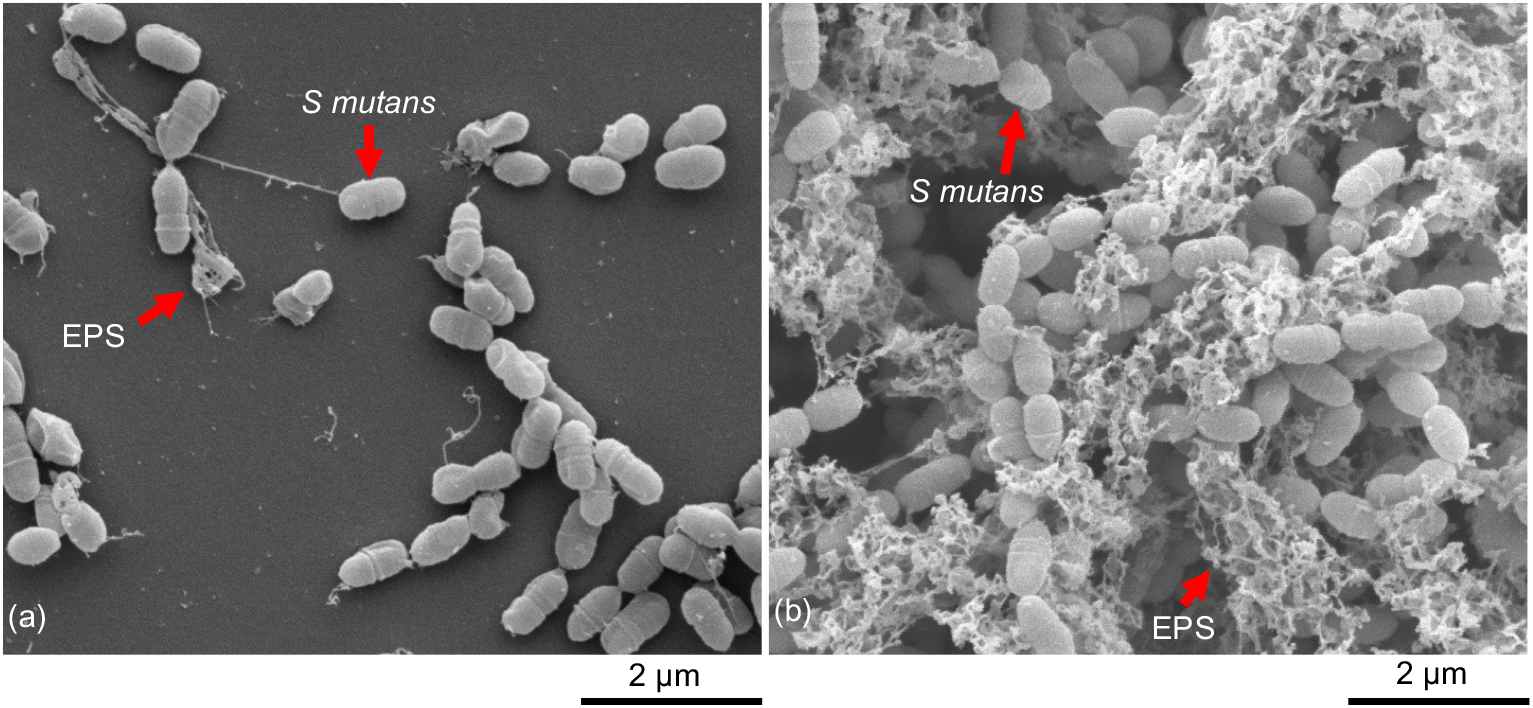
SEM images of *S. mutans* biofilms at 15000x magnification grown in THY on a titanium-coated substrate with (a) 0 mM sucrose (b) 75 mM sucrose.

In contrast, a biofilm grown with 75 mM sucrose has a thicker biofilm which covers the titanium surface of the substrate. More bacteria are present in the biofilm samples grown with sucrose, and the bacteria form longer chains of cells relative to the biofilms without sucrose. The increased thickness and coverage seen in the biofilms grown with sucrose agrees with previous studies showing increased thickness of biofilms with sucrose relative to biofilms grown in media without sucrose [36]. Biofilms with sucrose also show more EPS surrounding the cells relative to biofilms without sucrose (**Figure 3**(b)). EPS in biofilms grown with sucrose is plentiful and continuous across the imaged regions, rather than forming small distinct regions as in the biofilms without sucrose. Bacteria chains and the regions of varied coverage are also seen in previously published electron microscopy images of *S. mutans* biofilms [37, 38], though these studies focus on imaging techniques and substrate characteristics rather than the effect of nutrient concentration.

Biofilms grown with 0 mM sucrose (**Figure 4**(a)) have fewer cells and less EPS than biofilms grown with sucrose. Biofilms grown with 37.5, 75, and 375 mM sucrose have more complete *S. mutans* coverage, as visible in **Figure 4**(b), (c), and (d). The biofilms in these images are thicker than the biofilms without sucrose and show cells that have formed chains with abundant EPS. All biofilms at these concentrations have complete substrate coverage and no differences are observed for biofilm structure, biofilm quantity, or EPS formation at these concentrations. Though the concentration of sucrose differs across an order of magnitude for these biofilms, visual observation is insufficient to determine the effect of sucrose on biofilm formation. Biofilm presence in these images indicates that the viable sucrose concentration for biofilm growth has a wide range.

**Figure 4:**
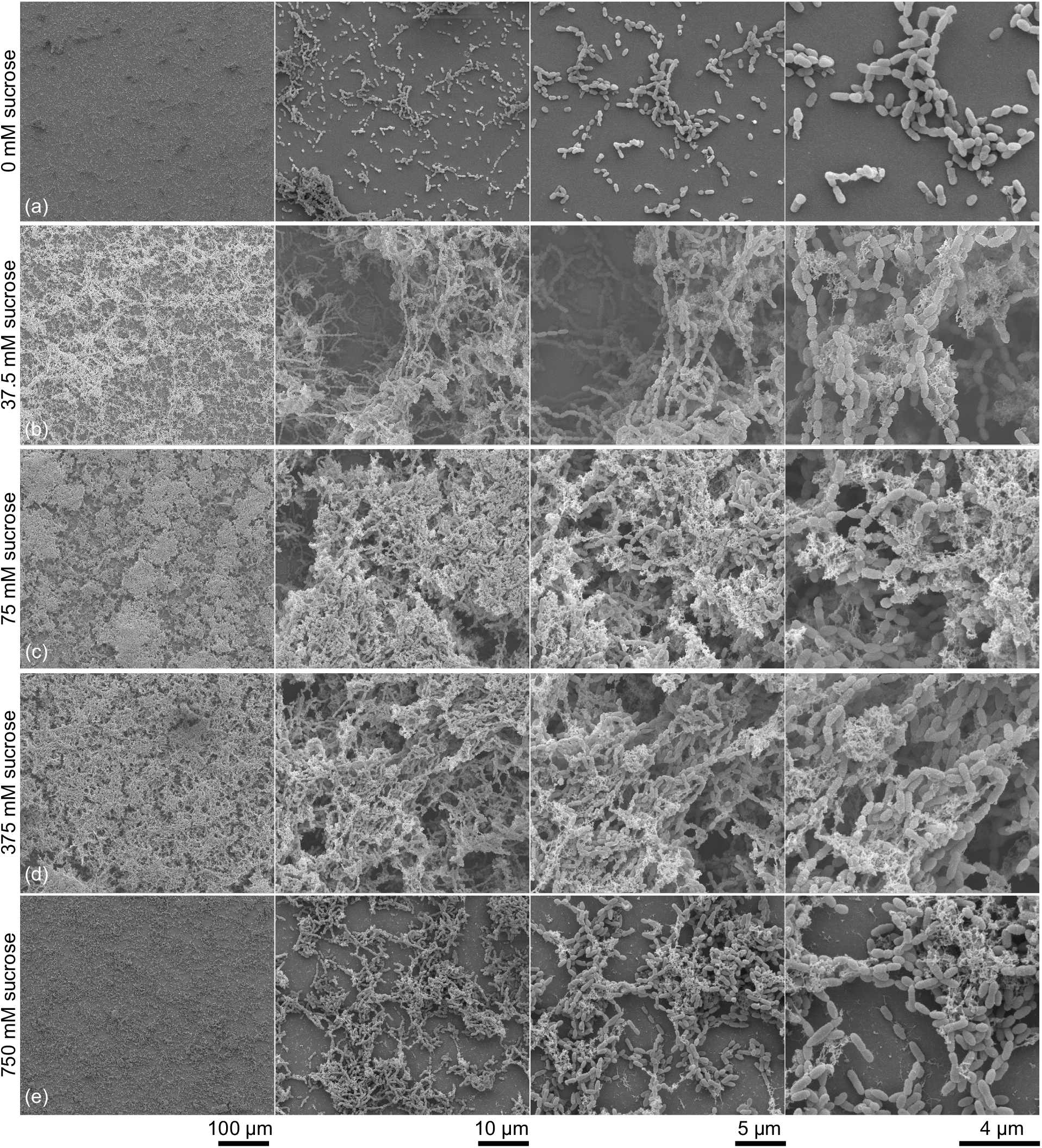
SEM images of *S. mutans* biofilms grown on titanium coated substrates with increasing concentrations of sucrose by row and increasing magnification (Left to Right: 250x, 2500x, 5000x, 10000x) (a) 0 mM sucrose (b) 37.5 mM sucrose (c) 75 mM sucrose (d) 375 mM sucrose (e) 750 mM.

At 750 mM sucrose, the biofilms have longer chains and increased surface coverage relative to the biofilms without sucrose, but have fewer cells and less EPS compared to biofilms grown with lesser concentrations of sucrose. In **Figure 4**(e), the titanium substrate can be seen underneath the 750 mM sucrose biofilm. Cai *et al*. demonstrate the effects of sucrose concentration on the volume of EPS in *S. mutans* biofilms [11, 39]. Biofilms in these studies that are grown without sucrose show no EPS. The authors also show that thickness of the biofilms and volume of the EPS initially increases with increasing sucrose, followed by a decrease in the thickness of the biofilm and the quantity of EPS above a certain concentration of sucrose, results that correlate with the results of the electron microscopy imaging presented here.

### 3.3 Electron Microscopy Profile Imaging

Top-down SEM imaging of *S. mutans* biofilms [30, 37, 40, 41] and biofilms of other species [42, 43] exists in the literature, and while this technique is useful for determining biofilm coverage and presence, top-down imaging cannot provide information about the height and formation of elevated features in the biofilm.

Electron microscopy imaging of biofilm profiles reveals details about the structure and formation of biofilms for each sucrose concentration at high magnifications. The electron microscope is focused first on the edge of the substrate and then subsequently farther away to image elevated structures within the biofilms. **Figure 5** demonstrates the imaging process on a biofilm with 37.5 mM sucrose. The long depth of focus and variable working distance of an electron microscope enables imaging of complex biofilm features away from the edge of the substrate.

**Figure 5:**
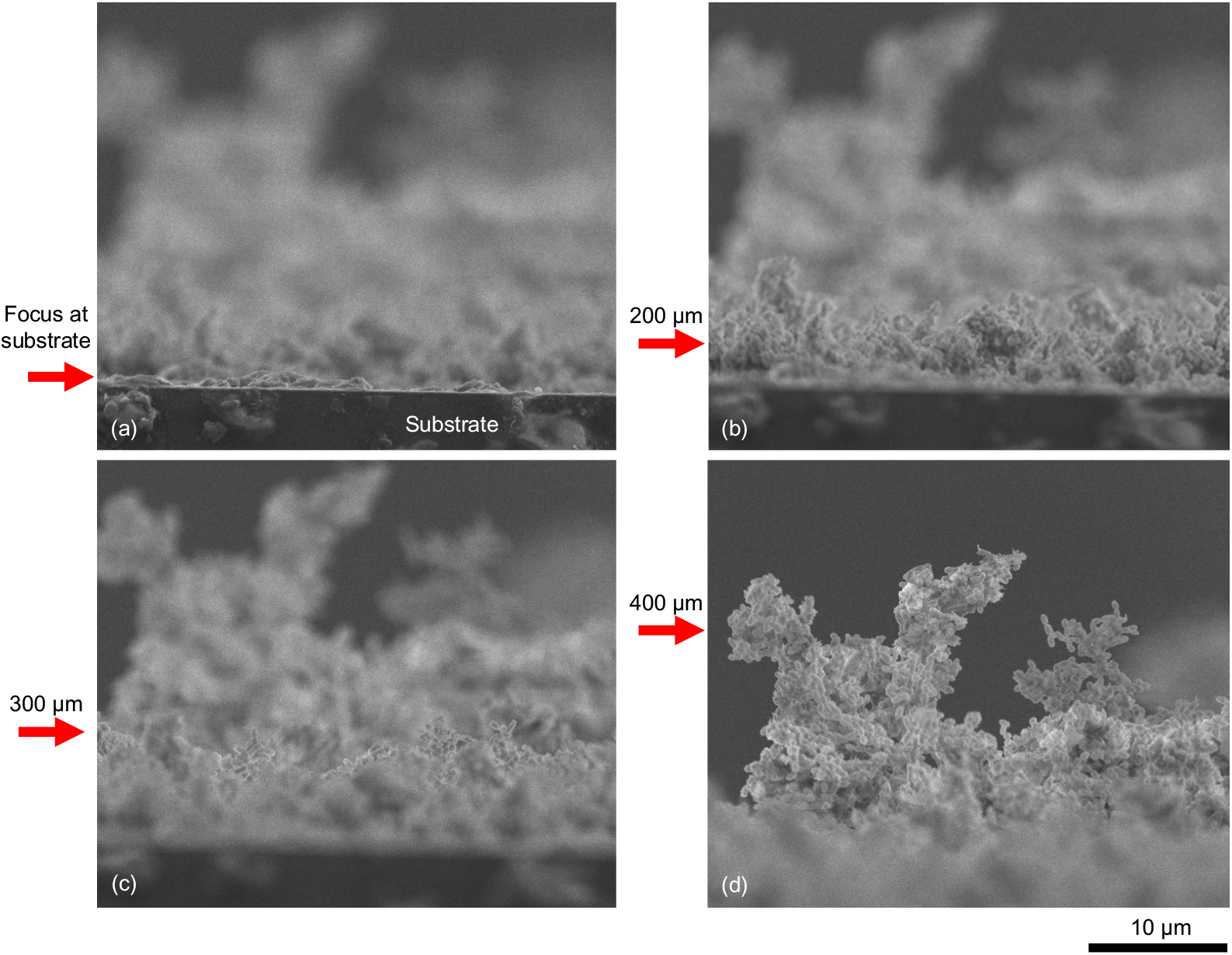
Profile SEM images of *S. mutans* biofilm with 37.5 mM sucrose on titanium coated substrates with arrows noting focused region at (a) edge of substrate (b) 200 μm from edge of substrate (c) 300 μm from edge of substrate (d) 400 μm from edge of substrate.

SEM profile imaging reveals that biofilms do not have a consistent profile or thickness at this scale, but instead have intermittent elevated features with complex geometries. These elevated features could be seen in top-down imaging, but specific details of the features are more easily seen in profile imaging. The elevated features are often delicate, as seen in **Figure 5** and **Figure 6**, and consist of both bacterial cells and EPS. Images from a biofilm grown with 37.5 mM sucrose show a large cluster of cells cantilevered from a narrow support only a few cells across (**Figure 6**(a)) and a single chain of cells which extends vertically surrounded by EPS (**Figure 6**(b)). These fragile features, which cannot be discerned with top-down SEM imaging, demonstrate the effectiveness of both the fixation method and the SEM technique for imaging biofilm formation.

**Figure 6:**
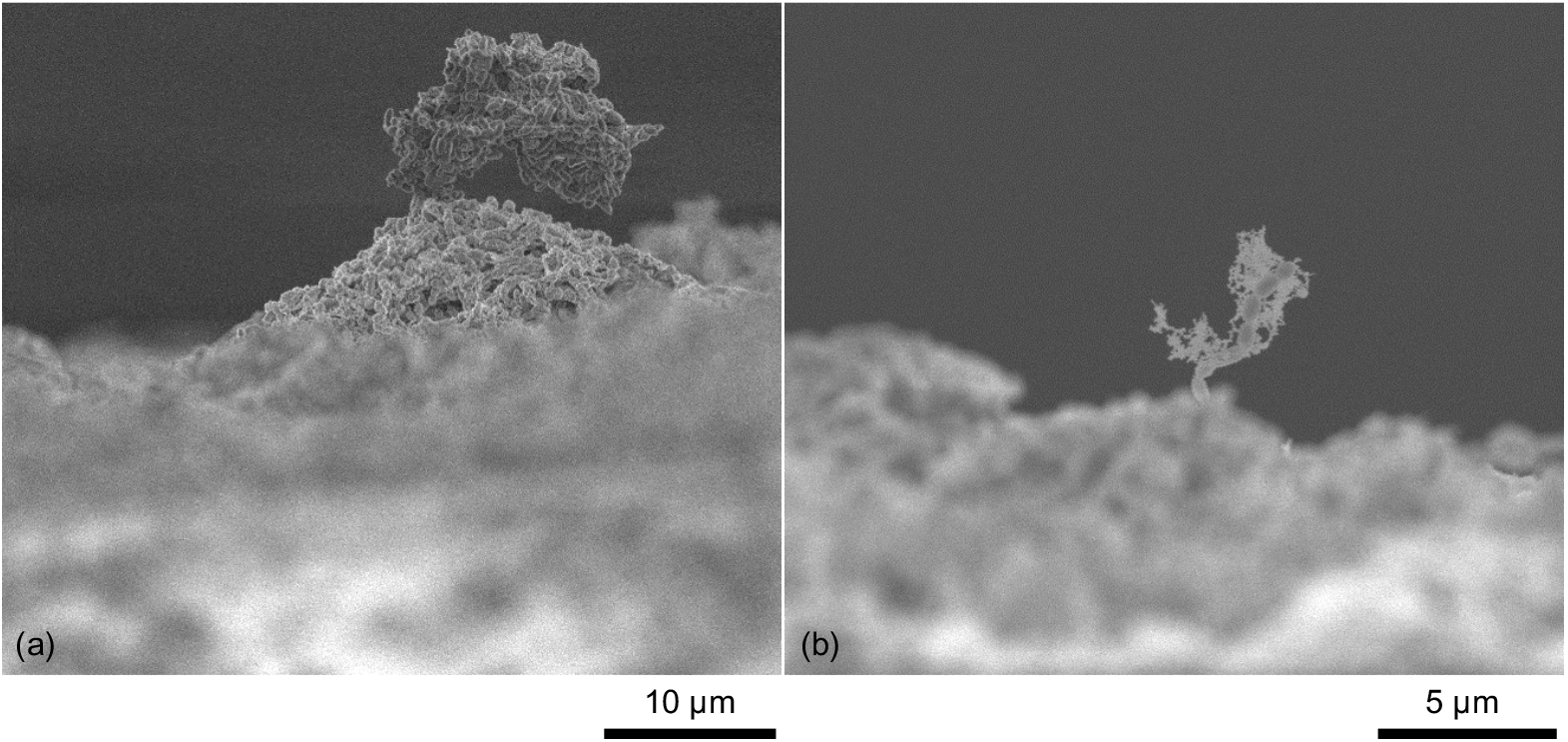
SEM images of structures in a *S. mutans* biofilm grown with 37.5 mM sucrose demonstrating that the microscopy preparation technique preserves (a) the formation of bacteria cells and (b) EPS formation.

In contrast, biofilms grown without sucrose do not exhibit complete substrate coverage or complex features seen in all biofilms grown with sucrose in this study. Profile imaging shows that the biofilm, where present, is commonly only a single cell in thickness. Small, sparse mounds that are clusters of cells greater than one cell in thickness are seen in 83% of locations imaged on sucrose-free biofilms during this study. All imaged mounds are less than 10 μm tall, as seen in **Figure 7**. Profile images of the 0 mM sucrose biofilms agree with both optical and top-down SEM images of biofilms in this study, which show that biofilms grown without sucrose have much less substrate coverage than those grown with sucrose.

**Figure 7:**
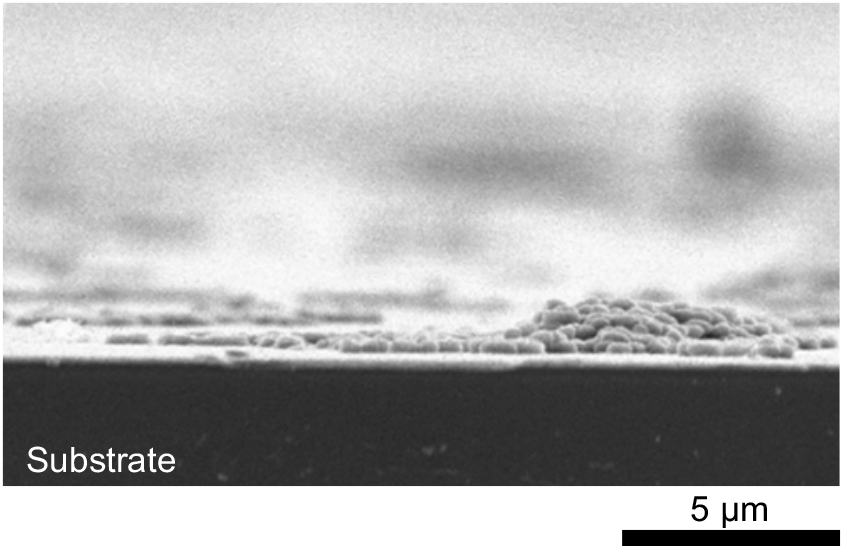
SEM profile image of *S. mutans* biofilm grown on a titanium-coated substrate with 0 mM sucrose.

All biofilms grown with sucrose have sufficient coverage such that the substrates could not be seen in profile imaging. Biofilms grown with sucrose also have mounds of bacteria and EPS that are distinct and elevated above the more consistent lower regions of the biofilm. Biofilms with 75 or 375 mM sucrose have thicker substrate coverage and higher elevated mounds than biofilms with either 37.5 or 750 mM sucrose. Biofilms with 75 and 375 mM sucrose each have multiple mounds above 100 μm in height, but the maximum feature height for 37.5 or 750 mM biofilms is less than 50 μm. Elevated mounds in biofilms with 75 and 375 mM sucrose are shown in **Figure 8**. The presence of large mounds in biofilms grown with both 75 and 375 mM sucrose indicates a large range of sucrose concentrations is viable for biofilm growth. In a previous study of biofilm formation using confocal laser scanning microscopy, Cai *et al*. described the presence of similar bacterial mounds in *S. mutans* biofilms, as well as increased quantities of EPS in biofilms between 30-300 mM sucrose relative to biofilms with 0 or 1100 mM sucrose [39].

**Figure 8:**
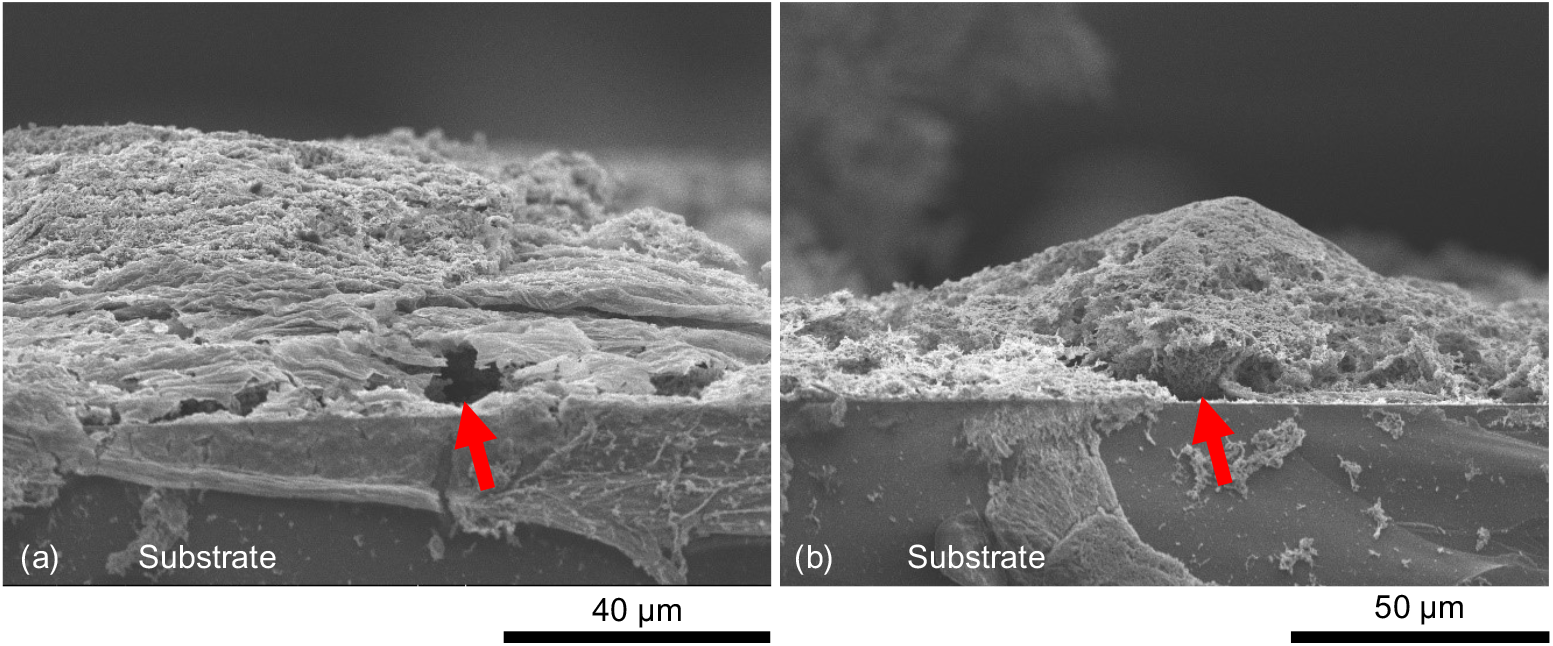
SEM images of channels inside thick regions of *S. mutans* biofilm grown with (a) 75 mM sucrose (b) 375 mM sucrose. Multiple channel-like regions are visible at the base of the biofilm mounds, one of which is indicated by the arrow.

Hollow areas at the base of elevated mounds are visible when the bases were not blocked by other features. Bacterial biofilms form in “microcolonies” consisting of bacteria, EPS, and channels for fluid transport [8, 44]. Hollow features noted in **Figure 8** could be these channels, and similar features are reported in other studies using top-down SEM imaging [37].

### 3.4 Laser Spallation

Weibull modeling of film failure is shown in **Figure 9**. Alpha and beta, the Weibull parameters used to produce the curves in **Figure 9**, and the adhesion strength values for each sucrose concentration are included in **Table 1**. Adhesion results indicate that sucrose plays a key role in the adhesion strength of *S. mutans* biofilms and that these biofilms are tolerant of a wide range of sucrose concentrations. The relationship between sucrose concentration and adhesion strength suggests an optimal concentration around 75 mM that maximizes the adhesion of *S. mutans* biofilms to titanium.

**Table 1:** Adhesion strength for each sucrose concentration, corresponding Weibull parameters, and root mean square (RMS) difference between Weibull model and experimental data. Percentile bootstrap estimates are used to produce the 95% confidence intervals listed in parenthesis. *There is no difference between model fit and experimental data.

| Sucrose (mM) | Adhesion Strength (MPa) | $\alpha$ Parameter | $\beta$ Parameter | RMS |
| --- | --- | --- | --- | --- |
| 0 | 89.1<br>(77.8, 100.4) | 92.7<br>(80.6, 104.8) | 9.38<br>(2.9, 29.9) | 0.0000* |
| 37.5 | 167.3<br>(157.0, 178.3) | 172.4<br>(163.2, 182.1) | 12.2<br>(4.3, 34.6) | 0.0000* |
| 75 | 336<br>(329.5, 342.5) | 337.9<br>(329.8, 346.0) | 66.6<br>(35.6, 124.5) | 0.0433 |
| 375 | 301.5<br>(291.9, 311.5) | 310.9<br>(300.5, 321.7) | 12.0<br>(3.7, 39.0) | 0.0383 |
| 750 | 262.9<br>(247.8, 278.0) | 267.1<br>(251.5, 282.7) | 23.53<br>(11.8, 46.8) | 0.367 |

**Figure 9:**
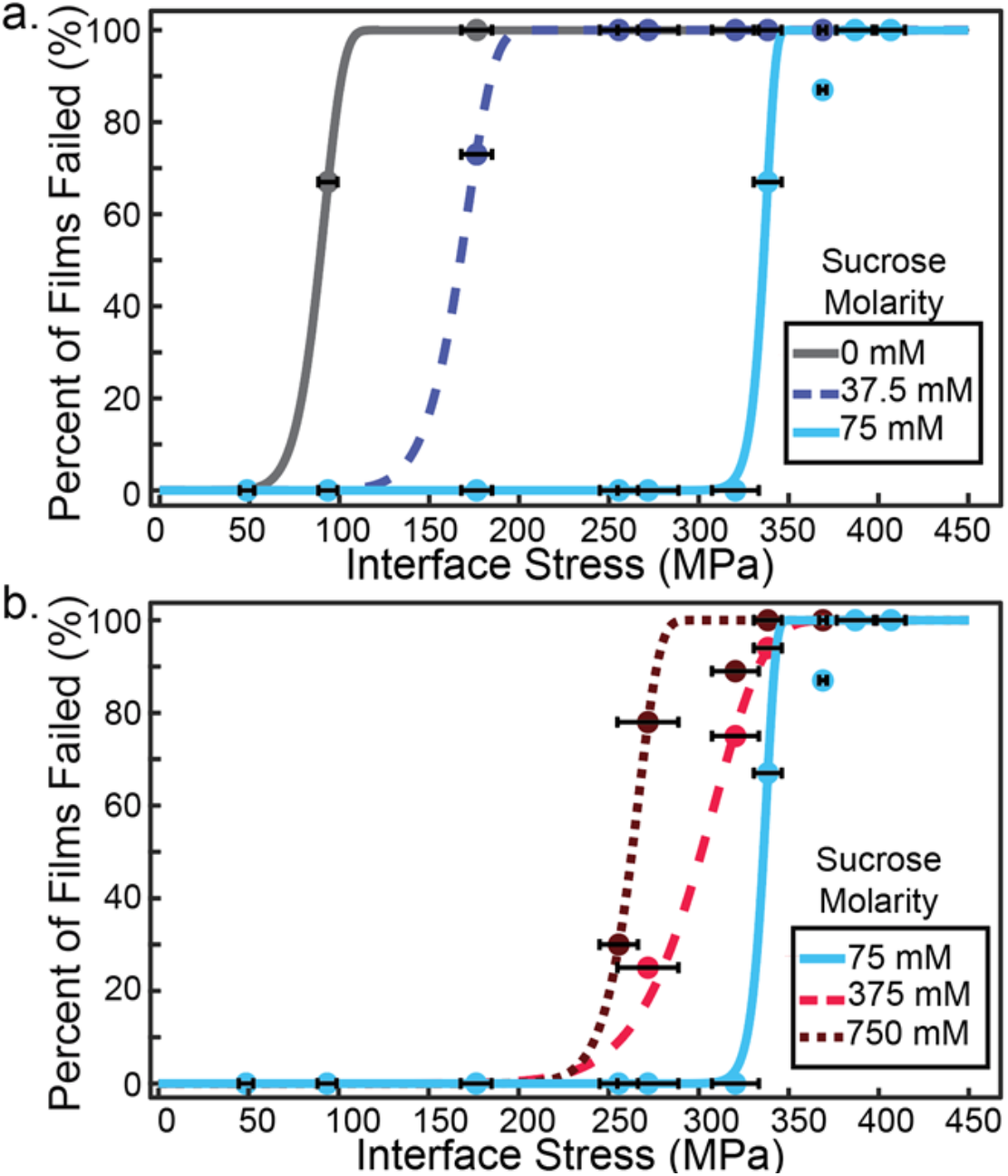
Percent of films failed when loaded with increasing interface stress. Data points represent experimental data. Lines represent the Weibull interpolation of experimental data. (a) Biofilms cultured with sucrose concentrations of 0-75 mM present an increase in adhesion strength with increasing sucrose concentrations. (b) Biofilms cultured with sucrose concentrations of 75-750 mM present a decrease in adhesion strength with increasing sucrose concentrations. Horizontal error bars are the standard deviation of the calibrated interface stress at each point.

For biofilms grown with 0 mM sucrose, the measured adhesion strength is 89.1 MPa with a 95% CI (77.8, 100.4). The adhesion strength increases for biofilms grown with 37.5 mM sucrose which have a measured adhesion strength of 167.3 MPa with 95% CI (157.0, 178.3). Biofilms grown with 75 mM sucrose exhibit the maximum measured adhesion strength of tested concentrations at 336 MPa with a 95% CI (329.5, 342.5). Above 75 mM sucrose, the measured adhesion strength decreases with increasing sucrose concentration. Biofilms grown with 375 mM sucrose exhibit a measured adhesion strength of 301.5 MPa with 95% CI (291.9, 311.5). Biofilms with 750 mM sucrose, the maximum concentration tested in this study, exhibit a measured adhesion strength of 292.9 MPa and a 95% CI (247.8, 278.0).

While a statistical decrease in adhesion strength is seen for sucrose concentrations above 75 mM, the decrease is not large relative to the increase in sucrose concentration. Between 75 mM and 750 mM sucrose, the decrease in adhesion strength is 22% while the sucrose concentration increases by an order of magnitude. Adhesion strength results supports the trend seen in this work that a wide range of sucrose concentrations exists that is viable for biofilm growth. Differences in biofilm characteristics for individual sucrose concentrations are small relative to the change in magnitude in sucrose concentration, a trend which has been seen in other studies quantifying the amount of EPS in *S. mutans* biofilms [39].

Adhesion strength values of all *S. mutans* biofilms in this study are shown in **Figure 10** and compared to measured adhesion strength of osteoblast-like cells from Boyd *et al*. [19]. Boyd *et al*. used the laser spallation technique to measure the adhesion strength of MG63 osteosarcoma cells to titanium coated substrates and found their adhesion strength to be 143 MPa with 95% CI (114, 176) [19]. Without sucrose, the measured adhesion strength of *S. mutans* biofilms is less than that of the osteoblast-mimicking cells, so the body may have a better chance to resolve *S. mutans* colonies. However, in the presence of 37.5 mM sucrose, and the higher sucrose concentrations included in this study, the lowest sucrose concentration tested in this work, the adhesion strength of *S. mutans* biofilms becomes stronger than the adhesion strength of MG 63 cells.

**Figure 10:**
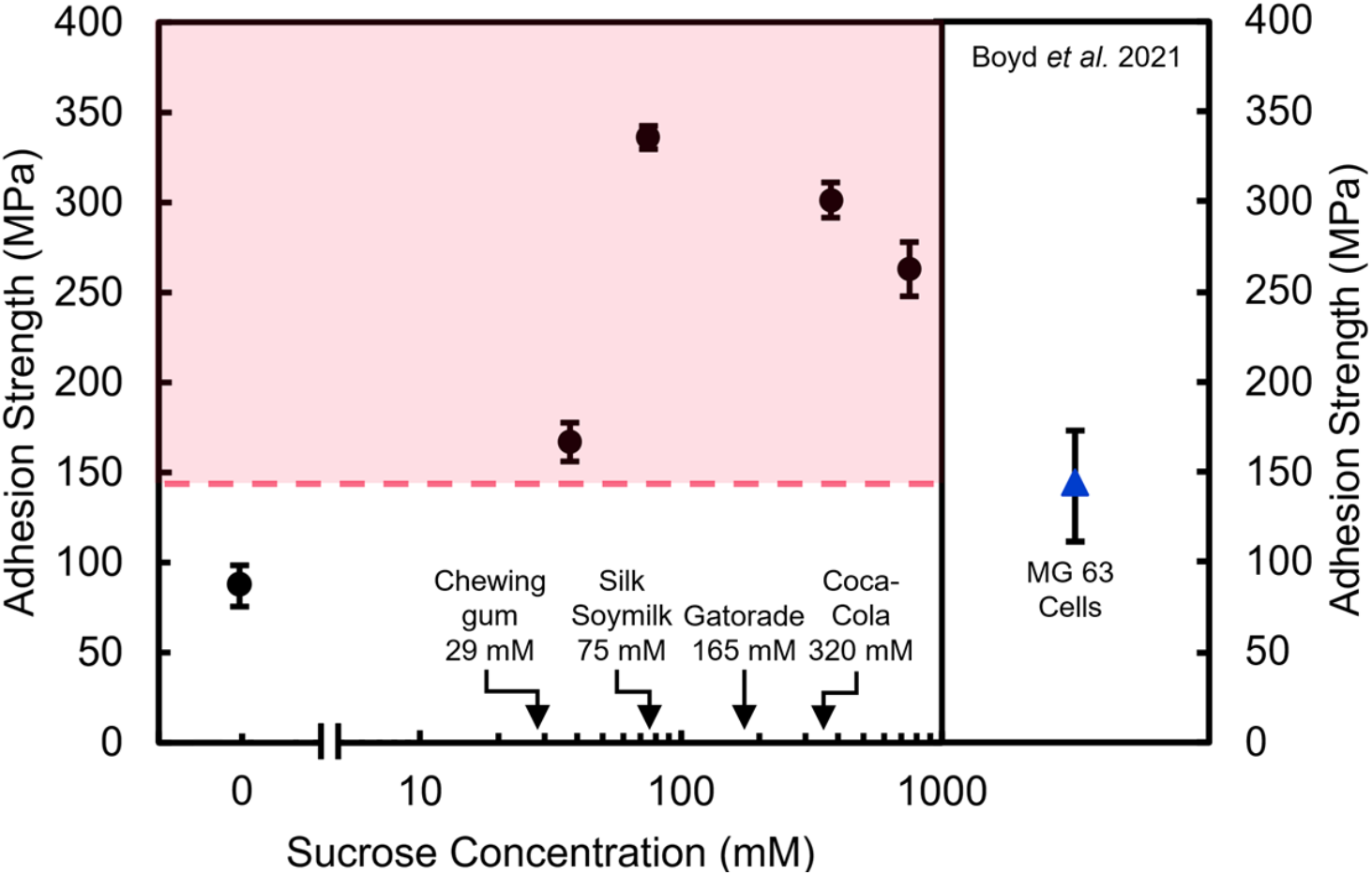
Black circles represent *S. mutans* biofilm adhesion strength to titanium substrates for each experimental sucrose concentration. The blue triangle and dashed red line show adhesion strength of MG 63 cells to the same substrates as previously determined by Boyd *et al*. [19]. Error bars are the 95% CI determined by percentile bootstrap estimates. The sucrose concentrations of common commercial food products are noted [45-48].

The range of sucrose concentrations in this work is relevant to common food items, which increase sucrose in the environment surrounding a dental implant during consumption. A previous study by Dawes *et al*. [45] determined that the concentration of sucrose in saliva after 10 min of gum chewing is approximately 30 mM. A commercial soy milk has a sucrose concentration of approximately 73 mM [46], similar to the sucrose concentration associated with the maximum biofilm adhesion strength found in this work. Lemon-lime Gatorade has a sucrose concentration of 170 mM [47]. Coca-Cola has a sucrose concentration of approximately 320 mM [48]. Other common foods have still higher sucrose concentrations, such as a commercial fruit jam which has a sucrose concentration of over 2 M [49].

Diets high in sugar are known to contribute to the risk of the development of dental caries [50]. Initial studies into the relationship between sugar and peri-implantitis show that sugar consumption has a plaque-promoting effect at implant sites as well as an association with peri-implant mucositis and peri-implantitis [51]. Experimental animal studies have also used processed, high-carbohydrate diets to provoke corresponding peri-implant inflammation [52]. Sucrose present in the implant environment that contributes to prolific biofilm growth and increased biofilm adhesion to the implant surface likely increases the risk of developing peri-implantitis as well.

## 4. CONCLUSIONS

Robust and adherent biofilm colonization on an implant surface can lead to infection. The environment surrounding dental implants including substrate material and nutrient concentration, affect the propensity for biofilm growth and thus may alter the propensity for infections to occur. In this work, the adhesion and formation of *S. mutans* biofilms on titanium substrates are studied in the presence of varying sucrose concentration from 0 mM to 750 mM. Biofilms grown with 0 mM sucrose form much shorter chains and much less EPS than biofilms grown with sucrose, and the sporadic substrate coverage can be seen optically. In biofilms grown with sucrose, elevated mounds and fragile features comprised of cells and EPS are preserved and imaged with electron microscopy. The highest elevated mounds are seen in biofilms grown with 75 and 375 mM sucrose and exhibit transport channels as reported in other work. A wide range of sucrose concentrations resulted in plentiful formation and adhesion of *S. mutans* biofilms on implant surfaces.

The laser spallation method is used to quantitatively determine the adhesion strength of the biofilms to titanium substrates. Maximum adhesion strength occurs in biofilms grown with 75 mM sucrose. For sucrose concentrations above 75 mM, the adhesion strength decreases, and biofilms grown with 750 mM sucrose have an adhesion strength that is 22% lower than 75 mM sucrose. The decrease in adhesion strength is small relative to the increase in sucrose, which agrees with trends seen in electron microscopy images of the same biofilms. The condition that produced the least adherent biofilm is the growth condition that lacked sucrose altogether and registered an adhesion strength that is 73% lower than the 75 mM sucrose condition. Sucrose enhanced the EPS production of *S. mutans* biofilms and increased the adhesion strength to titanium. Adhesion strength of *S. mutans* biofilms with sucrose to titanium is greater than the adhesion strength of osteoblast-like cells to the same surface, which suggests that sugary foods may help *S. mutans* biofilms outcompete osteoblasts during osseointegration. Sugar-rich diets are a well-known risk factor for dental caries and are also associated with peri-implantitis. Sucrose-mediated biofilm adhesion and formation on titanium is a possible mechanism to explain how sucrose in the environment could lead to higher rates of peri-implantitis.

## ACKNOWLEDGMENTS

Scanning electron microscopy was performed in the Electron Microscopy Center at the University of Kentucky, and the authors wish to thank Nicolas Briot for assistance with specimen preparation. The authors wish to thank Natalia Korotkova for the generous donation of *S. mutans Xc* and Arnold Stromberg with the Applied Statistics Laboratory for the Weibull modelling and analysis. The authors also wish to thank Dr. Craig Miller and Dr. David DeVito for their guidance.

## AUTHOR CONTRIBUTIONS

Methodology, all authors; formal analysis, L.W. and J.B.; investigation, L.W., T.B., and J.B.; data curation, T.B.; writing—original draft preparation, all authors.; writing—review and editing, L.W. and M.G.; supervision, M.G.; funding acquisition, M.G. All authors have read and agreed to the published version of the manuscript.

## DATA AVAILABILITY STATEMENT

The raw data required to reproduce these findings are available to download from https://doi.org/10.18126/d1bg-nwtg via the Materials Data Facility [53, 54].

